# Effects of gamebird release on the non-gamebird dietary diversity and composition of red foxes (*Vulpes vulpes*) in Southern England

**DOI:** 10.64898/2026.08.20.745779

**Authors:** Siffreya Pedersen, Rufus B. Sage, Maureen I. A. Woodburn, Jenny R. Coomes, Joseph Werling, Charles R. Tyler

## Abstract

The red fox, *Vulpes vulpes*, is abundant in England and can exert limiting effects on their avian and mammalian prey. Large-scale gamebird releases in the UK may be sustaining high predator numbers, leading to greater predation on other prey species, especially once gamebird stocks are depleted. However, little is known how gamebird release affects the broader diet of the red fox. Here we used DNA metabarcoding to assess the diet of foxes from 18 agricultural estates in Southern England, 10 of which released large numbers of gamebirds (red-legged partridges – *Alectoris rufus*, and common pheasants – *Phasianus colchicus*) and 8 which did not. Scats were collected over one year, allowing for seasonal investigation of the foxes’ diet. We investigated the vertebrate species consumed and compared the non-gamebird dietary diversity and composition between release and non-release estates and across seasons. The field vole (*Microtus agrestis*) was found to be the most frequently predated species overall. Brown hares and field voles were detected more on release sites, while bank voles and dog faeces were detected more on non-release sites. We found little evidence that foxes predate ground nesting birds or other species of concern. The dietary diversity was significantly lower on estates that released gamebirds, and this difference was most notable during the post-shoot, spring months (February to April). On both estate types, diversity was highest in the summer months. The altered predatory behaviour due to gamebird release is likely to affect the populations of the non-game prey of the red fox and this should be considered when designing policies regarding gamebird management and biodiversity conservation. Alternative predation combined with predator control may be reducing predation pressure on non-game prey where gamebirds are released, but high fox density elsewhere likely results in higher predation pressure on a wide range of species.

## 1. Introduction

An estimated 32 million common pheasants (*Phasianus colchicus*) and 9 million red-legged partridges (*Alectoris rufus*) are released annually into the UK landscape to support recreational hunting (Madden, 2021). It has been estimated that at the time of their release in late summer, these two species comprise ∼ 45 % of the British bird biomass, with this figure dropping to ∼ 12 % in spring (Blackburn & Gaston, 2021). The scale of gamebird release has prompted interest in their ecological impacts (Madden et al., 2023; Madden & Sage, 2020; Mason et al., 2020; Sage et al., 2020). One particular concern is the effect that gamebird release could have on the abundances of generalist predators, such as the red fox (*Vulpes vulpes*), by providing easy prey or carrion to feed on, and the implications of this on the wider ecology of these landscapes. Approximately 60 % of gamebirds released are not shot (Madden et al., 2018), and after shooting, the most common cause of death is via predation with approximately 32 % of the pheasants released ultimately being predated or scavenged by predators, primarily foxes (Madden et al., 2018; Turner, 2007). Gamebirds are released from June to August and predation by foxes occurs through the winter and into spring, before reducing in summer when gamebird availability is much lower (Madden et al., 2018; Sage et al., in prep.; Sage et al., 2025). The indirect effects of this on the other prey species is not well understood. On one hand, gamebird release may benefit prey species by diverting predation pressure away from them (Reshamwala et al., 2018; Rodewald et al., 2011). Conversely, it has been hypothesised that gamebird releases may increase the abundance of foxes, and that this could result in increased predation on non-game species once gamebird numbers have declined (Madden & Sage, 2020; Mason et al., 2020).

The species that could be indirectly affected by change in fox predation or abundance are numerous. Studies into the diet of foxes in Britain (Baker et al., 2006; Doncaster et al., 1990; Lever, 1959; Reynolds et al., 1995; Reynolds & Tapper, 1995; Southern & Watson, 1941; Waggershauser et al., 2022; Webbon et al., 2006) and throughout the world (Castañeda et al., 2022) have found small mammals to be the foxes’ main prey group, followed by invertebrates, lagomorphs, fruit, birds, vegetation, large mammals, anthropogenic food, and lastly, reptiles. As generalist predators, their diet varies substantially dependent on the region and local availability of prey (O’Mahony et al., 1999; Reynolds et al., 1995; Soe et al., 2017). On arable estates in Southern England, lagomorphs have been found to be the primary prey species, followed by small mammals (Baker et al., 2006; Doncaster et al., 1990; Lever, 1959; Reynolds et al., 1995; Southern & Watson, 1941; Webbon et al., 2006). Despite the many studies into the diet of this ubiquitous predator, and how it varies depending on anthropogenic factors (Reshamwala et al., 2018; Williams et al., 2024) and the ecology of prey species (O’Mahony et al., 1999), no attempt has been made to investigate how the diet of foxes varies with gamebird release.

Here we investigate how gamebird release affects the non-game component of the diet of the red fox and attempt to contextualise this in terms of the wider ecology. To do this, we applied DNA metabarcoding on fox scats collected over a period of one year from agricultural estates in Southern England, some of which release large numbers of gamebirds. We assess the diversity and abundance of non-game vertebrate species detected in the scats and compare this across the seasons and between gamebird release and non-release sites.

## 2 Methods

### 2.1 Study sites and sample collection

The sites selected for study were all in Southern England, spread across the counties of Wiltshire, Southern Hampshire, and Northeastern Dorset. Gamebirds, specifically pheasants and red-legged partridges, were released in moderate to large numbers (between 10,000 and 25,000 birds annually) on 10 of the sites, while another 8 sites had no gamebird release (see Sage et al., 2025 for further site details). The sites were not paired but selected to be as similar as possible in land use and were all predominantly arable with a smaller amount of grassland, interspersed with areas of broadleaf woodland (Figure 1, section 2.4).

**Figure 1.**
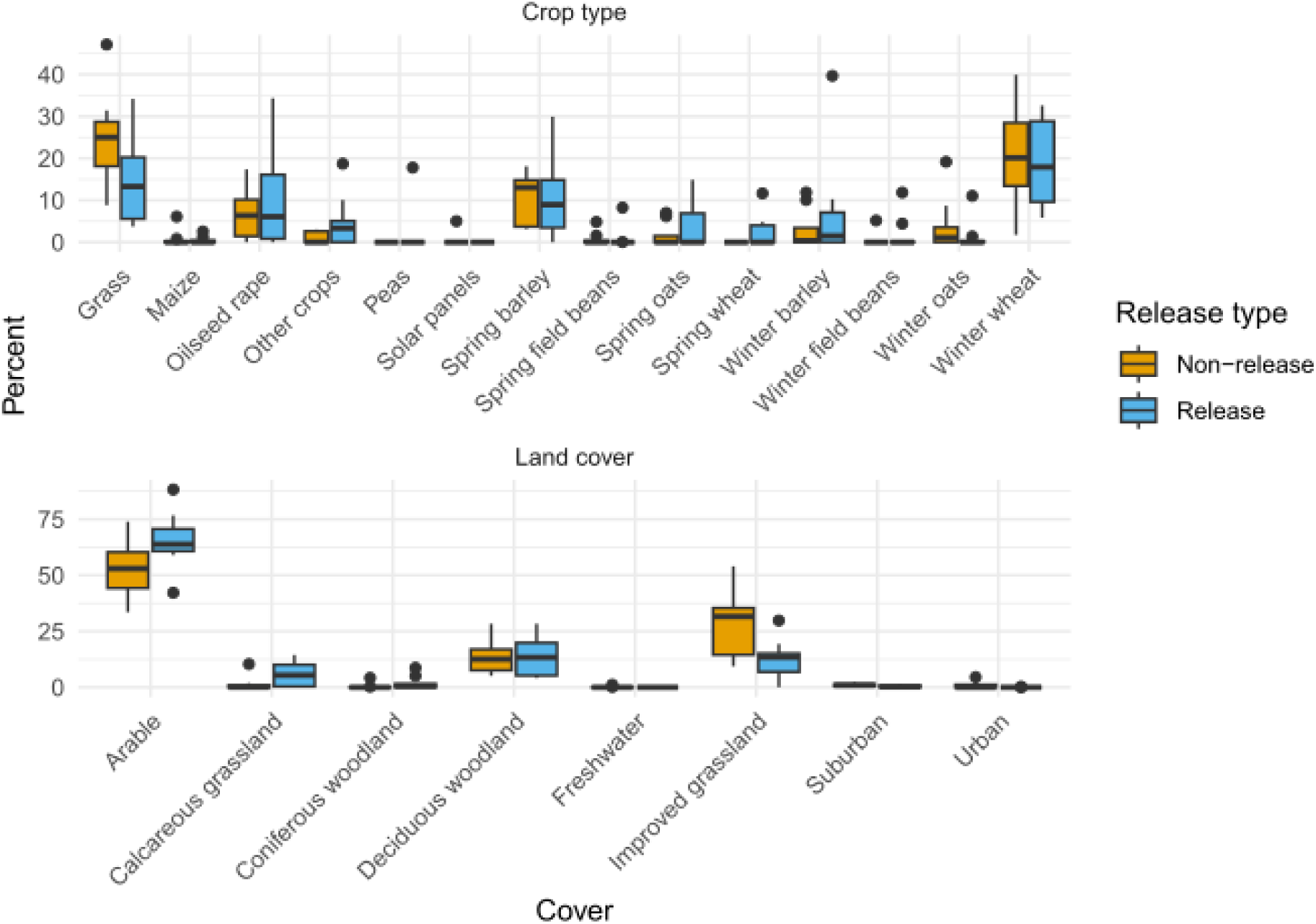
Land cover and crop types within 1 km from each site. Percentage cover within 1 km of the centre of each site transect was calculated from UKCEH 2022 Land Cover and 2022 Land Cover plus: Crop maps.

Additionally, the average distance to an urban area was similar between gamebird release and non-release sites (Sage et al., 2025). Some spread of gamebirds into non-release sites was expected as 3 non-release sites shared a boundary with a release site, and the other 5 had boundaries within 1km. Finding sites far from release pens with comparable habitat and within the same region was not possible due to the high rate of gamebird release in Southern England (Madden & Sage, 2020). However, bird counts were conducted at the same time as fox scat collection and the gamebird abundances were found to be considerably higher on release sites compared to non-release sites (Sage et al., 2025).

Details of scat sample collection can be found in Sage et al. (in prep.). Briefly, a fixed route of approximately 3 km was walked every 3 weeks on each site, from February 2022 to March 2023. Fox scats were identified based on appearance and scent and were collected and frozen on the same day. Molecular analysis was used to confirm the scat identity being fox (see section 2.3.3).

### 2.2 Molecular analysis

#### 2.2.1 Subsampling

Approximately 0.5 cm^3^ of each scat was taken for DNA extraction. Equal volumes, rather than weights, were used to regulate the amount of material extracted from as the samples differed in their water content. To obtain a representative sample, 3 to 5 subsamples were taken from along the length of the scat. Wherever possible, the subsamples were taken from inside the scat to reduce chances of environmental contamination. The forceps and spatula used for sampling were sterilised in 10 % bleach solution, rinsed in deionised water, and rinsed again in 70 % ethanol between samples.

#### 2.2.2 Extraction

DNA was extracted with the QIAamp Fast DNA Stool Mini kit (Qiagen) following the protocol for isolation of DNA for human DNA analysis with the following modifications; samples were heated to 70 °C for 5 minutes in 1 ml InhibitEX buffer to increase the chance of lysis of hair and bone, and the DNA was eluted in 100 μl ATE buffer with 5 minutes incubation at room temperature to increase the final concentration. For every 29 fox scat sample DNA extractions a blank extraction was included. DNA extracts, including blank extracts, were transferred to 96 well plates ensuring that collection sites and months were randomly spread between plates to mitigate for any possible plate effects on the results. Extracts were quantified with the QuantiFluor ONE dsDNA System (Promega) and normalised to 1 ng/μl.

#### 2.2.3 Library prep

A ∼ 106 bp region of the 12SV5 gene region was amplified with primers as derived by Riaz et al. (2011) (F:5’-TTAGATACCCCACTATGC-3’, R:5-TAGAACAGGCTCCTCTAG-3’). To allow the samples to be pooled and sequenced in a single run, 96 pairs of primers were purchased each with 12 nt unique indexes (from Caporaso et al., 2012) on the 5’ end of both forward and reverse primers, with a CT spacer between index and primer. Indexes are short regions of DNA attached to the primers and are amplified along with the template DNA, thus allowing samples from the same plate to be pooled once amplified and separated bioinformatically later. PCR reactions consisted of 25 μl OneTaq 2x Master Mix with Standard Buffer, 0.2 μM of each primer, 5 ng template DNA, and 0.2 μM of the canid blocking primer RacBlk (Woo et al., 2022), made up to 50 μl with HPLC water. Thermocycling conditions started with an initial denaturation at 94 °C for 30 seconds, then 35 cycles of 94 °C for 30 seconds, 58 °C for 30 seconds, and 68 °C for 30 seconds, with a final extension at 68 °C for 5 minutes. PCR success was assessed by running each sample of a 2 % agarose gel.

Three no template controls were randomly distributed in each plate and one positive control. The no template controls were run as above but with no template DNA and were to control for contamination of the PCR and cross contamination between wells of the PCR plate. The positive control DNA consisted of 1 % pheasant DNA and 1 % partridge DNA in an equimolar mix of fox, jay, magpie, rat, grey squirrel, grass snake, adder, slowworm, and frog DNA that we derived from samples we collected directly from these animals and extracted with the DNeasy Blood & Tissue Kits (Qiagen). Animal samples were collected in accordance with the Animal Welfare Act 2006; snake (grass snake and adder) samples were taken from sloughed skins donated to us by the Amphibian and Reptile Conservation Trust, jay, slow worm, and frog samples from road-kills, and fox and magpie from legal predator control events.

The 12SV5 gene is both short and variable in length between species, and so the amplicons were cleaned with a 2x volume of SPRI beads to ensure amplicons from species with shorter 12SV5 genes were retained. Amplicons were quantified with the QuantiFluor ONE dsDNA System (Promega), and successful reactions from each plate were then pooled in an equimolar manner to give 7 pools (1 per plate of reactions). As the blank and no-template control reactions could not be quantified, 2 μl of each were added to their respective pool. The seven pools were further cleaned with a 2x volume of SPRI beads to ensure complete removal of residual primers and to concentrate each pool. A TapeStation (Agilent) using D1000 ScreenTape was used to confirm the complete removal of primers and to quantify the DNA concentration in the final pools. Illumina adapters and further unique dual indexes were ligated to each pool by the Exeter Sequencing Service, thus allowing all samples to be pooled together for sequencing. Sequencing was done on a NovaSeq 6000 by the Exeter Sequencing Service.

### 2.3 Bioinformatics

#### 2.3.1 Demultiplexing

Sequencing data were returned by the Exeter Sequencing Service as 14 files, 2 files for each of the 7 pools. These files were demultiplexed further with Cutadapt v3.5 (Martin, 2011) by providing the full primer-spacer-barcode sequences as anchored 5’ adaptors, in paired-end format with default error tolerance. Ligation-based library preparation is non-directional, as a result the R1 and R2 files produced by Exeter Sequencing Service contained an equal mix of forward and reverse reads. To reorient the reads and make them suitable for the DADA2 pipeline, demultiplexing was carried out in two rounds. In round 1, the paired-end input files and barcode mapping files were input in the standard manner and all un-trimmed reads saved to unknown_R1 and unknown_R2 files. In round 2 the input files were the unknown files from round 1, with unknown_R2 provided as the first and unknown_R1 provided as the second input. This procedure produced 4 files for each sample; R1 and R2 files from round 1 in standard orientation, and R1 and R2 files from round 2 in which the reads had been reorientated so that all forward reads were in R1 files and vice versa.

Quality filtering, trimming, denoising and chimera detection was undertaken using DADA2 v1.26.0 (Callahan et al., 2016) in R v4.2.0 (R Core Team, 2021). The error rates DADA2 calculates and uses for error correction can differ between the forward and reverse reads output by Illumina, so quality filtering, trimming, and error correction were run separately for each round of demultiplexed data. Cutadapt was used to remove primers at the 3’ end of reads that were read in the reverse complement due to the 150 bp read length being longer than the amplified 12SV5 region for some species. Reads that contained ambiguous nucleotides, and with a higher maximum expected error than 1 were removed, and all reads were truncated to 70 bp with DADA2’s filterAndTrim function. Default settings were used to establish the error rates (learnErrors function), denoise the reads (dada), and merge the paired end reads (mergePairs). Having taken both rounds of demultiplexed data through the above process, the two resulting sequence tables were merged, summing the read counts for identical ASVs (actual sequence variants). Chimeras were then removed using the removeBimeraDenovo function with the method set to ‘consensus’.

To obtain reads that are representative at the species level, and ease downstream work, the ASVs output by DADA2 were clustered into operational taxonomic units (OTUs) with Swarm v3.1.0 (Mahé et al., 2021) with *d* set to the default (1). Finally, spurious OTUs were merged with their parent OTUs with LULU (Frøslev et al., 2017).

#### 2.3.2 Taxonomic assignment

Taxonomy was assigned by comparing the OTUs to the GenBank nucleotide database (accessed on 29^th^ May 2024), using BLAST 2.13.0+ (Camacho et al., 2009) and using default parameters, followed by implementing the lowest common ancestor (LCA) approach in Megan v6.24.1. To improve the taxonomic resolution, taxonomic assignment was adjusted manually to remove the effect of hits with species not known to occur in the UK. Where OTUs were not assigned to species level, the UK species inventory, GBIF, and the BTO website were used to assess the occurrence of species in Southern England, and matches to species that do not occur, or are very rare visitors, in Southern England were disregarded. When only one match remained, the OTU was assigned to that species. When there was more than one UK match, but one had a perfect match (defined as total overlap with 100 % similarity), the OTU was assigned to the species with the perfect match. When there were no perfect matches, or there were multiple perfect matches, the OTU was assigned to the lowest common ancestor. One OTU that identified as *Alectoris chukar*, which is no longer released in the UK, was assumed to be a hybrid *A. rufa x chukar* and assigned to *A. rufa*.

#### 2.3.3 Data curation

To remove the effects of low levels of cross contamination and background noise, OTUs read counts that were less than 1 % of the reads from that sample were changed to zero. Samples that had less than 50,000 reads were discarded; this cut-off was chosen based on a comparison of read numbers from real samples and negative controls. Human reads, which occurred at low levels (mean relative read abundance of 0.07 %), were removed as they were considered to be either environmental contamination or contamination from the researchers who handled the samples.

Samples were checked as deriving from fox by comparing the read numbers that identified as fox, and other predators and scavengers that produce scats or pellets that could be mis-identified with a fox scat. Taken from the list of identified taxa, these species were: barn owl (*Tyto alba*), domestic dog (*Canis lupus*), Eurasian badger (*Meles meles*), buzzard (*Buteo buteo*), kestrel (*Falco tinnunculus*), mink/polecat (*Mustela spp.*), and domestic cat (*Felis catus*). Any samples that had fewer fox reads than any other predator were discarded.

Samples that had no reads from any predator, including fox, were retained, as it was assumed that the Canidae blocking primer effectively blocked all fox reads in these cases.

After the source check detailed above, fox reads were discarded. Samples that had no reads remaining after the removal of the fox were retained as these may have been from foxes that had fed only on plants or invertebrates.

### 2.4 Data analysis

To mitigate bias caused by primer bias, variable mitochondrial copy number, and variable rates of DNA degradation in the digestive system, read numbers were transformed to presence/absence. All analyses were carried out with R version 4.4.3 (R Core Team, 2021), metabarcoding data was manipulated with the R package phyloseq (McMurdie & Holmes, 2013) and plots created with ggplot2 (Wickham, 2016).

As this study focuses on the comparison of fox predation on non-game species between release and non-release estates, all gamebird detections were removed from the dataset, all of which were the common pheasant or red-legged partridge. The effect of gamebird release on the predation of these birds is explored in Sage et al. (in prep.).

#### 2.4.1 Alpha diversity metrics

To assess variations in dietary diversity in the face of variable sample numbers, the package iNEXT (Chao et al., 2014; Hsieh et al., 2024) was used to standardise sample numbers to equal coverage. To do this, iNEXT estimates the number of samples that would be necessary to record a standard percentage of the true prey community present and uses rarefaction to calculate diversity for sites that have more samples than necessary, and extrapolation for sites that have too few. This has been suggested to be an equitable way to compare sites of differing diversities, as more diverse sites require higher sample numbers to record an equivalent proportion of the diversity compared to a lower diversity site (Roswell et al., 2021). The coverage of each group was standardised to the default of iNEXT, which is the minimum coverage across all groups extrapolated to double that sample size (Hsieh et al., 2024). iNEXT was used to calculate and compare observed richness (q = 0), Hill-Shannon (q = 1), and Hill-Simpson (q = 2). These three metrics are on a common scale (units = species equivalents) but differ in the way they weight rare species (Roswell et al., 2021). Observed richness is the count of species with each species weighted equally, regardless of abundance. Hill–Simpson emphasises dominant species more strongly. Hill–Shannon balances the contributions of common and rare species and is intermediate between observed richness and Hill-Simpson.

To assess dietary diversity between gamebird release and non-release sites, all samples from each site were grouped and the frequency of each species per site was calculated. The function estimateD was then used to standardize to coverage and calculate diversity metrics as above. One release site was removed from this analysis due to having too few samples. These site level diversity metrics were then used in general linear models (see section 2.4.3).

To assess changes in dietary diversity over the year, the data was grouped into four seasons based very approximately on the timings of gamebird rearing, release, and shooting (see also Sage et al. 2025). These seasons were the post-shoot season from Feb to April inclusive, the summer - May to July when most gamebirds are being reared or only recently released, the post-release season from August to October when gamebirds are released but most (pheasant) shooting has not yet started, and winter from November to end of January during which time the majority of the shooting takes place. Due to low sample numbers during winter, samples from different sites were grouped together based on season and gamebird release site type. Diversity estimates for each group were calculated as above and plotted with 95 % confidence intervals calculated with 50 bootstraps.

#### 2.4.2 Land cover measures

As even small variations in land use and habitat availability could affect the availability of prey species, thereby potentially affecting the comparison of fox diet on sites with and without gamebird release, such effects were assessed and accounted for. Custom extents of the Land Cover Map 2022 and Land Cover plus: Crops (2022) were downloaded from the UK Centre for Ecology and Hydrology (UKCEH) as vector maps via Digimap (UKCEH, 2022b, 2022a). These were used to calculate percentages of land cover and crop type within 1 km of the centre of the transect walk for each site using ArcGIS Pro 3.1.2 (Esri, 2023). A radius of 1 km was chosen as this size would cover each transect entirely and was roughly similar to the average territory size of the red fox in rural Southern England (2.7 km2; Reynolds et al., 1995). This resulted in two datasets: one for land cover that including the percentages of arable land, calcareous grassland, coniferous woodland, deciduous woodland, freshwater, improved grassland, suburban and urban land within 1 km. The second dataset was for crop type and included percentages of grass, maize, oilseed rape, peas, solar panels, spring barley, spring field beans, spring oats, spring wheat, winter barley, winter field beans, winter oats, and winter wheat As the land cover and crop type data were compositional (i.e. relative parts constrained to a constant sum), they were centre log ratio transformed prior to use with the decostand function in vegan (Oksanen et al., 2022), opting for the ‘robust clr’ method to handle zeros. This step was necessary to remove the inherent negative correlations between components of such data and make it suitable for use in generalised linear models, which require independent, unconstrained covariates. Some variables were both very rare and strongly confounded with gamebird release or another variable and were removed prior to analysis, these were: freshwater from the land cover data, and peas, solar panels, spring wheat, and spring field beans from the crop type data (Figure 1). To facilitate interpretation and avoid nested predictors (i.e. crop types nested within arable land cover), the land cover and crop type data were incorporated into model selection separately.

#### 2.4.3 Statistical analysis

Generalised linear models were used to assess whether the three dietary diversity metrics were affected by gamebird release at the site level, while controlling for variations in land use. The models were fit with Gamma distribution with log link, after inspection of model diagnostic plots for each diversity metric with gamebird release treatment as the only predictor. The dredge function from the R package MuMIn was used to perform all subsets variable selection based on corrected AIC (AICc), with gamebird release treatment included in every subset via the argument ‘fixed’. This variable selection procedure was run 6 times in total, for each combination of the 3 diversity metrics and 2 land use datasets. The Benjamini & Hochberg method was used to control the false discovery rate and account for multiple testing.

Generalised linear mixed models were used to assess the effect of gamebird release and season on the composition of the diet, while controlling for variations in land use. GLMMs were fit with a multivariate response variable using the gllvm R package (Niku et al., 2019), with the number of latent variables set to 0, after model comparisons indicated this produced the lowest AIC. No sample grouping was necessary for this analysis, so the response variable was a scat x species presence/absence matrix, and the models therefore fit with a binomial distribution. Species that were found in fewer than 5% of scats were grouped: the Chinese muntjac, roe deer, fallow deer, wild boar, domestic cattle, sheep, and cat into ‘large mammals’; the Eastern grey squirrel, Eurasian pygmy shrew, Eurasian common shrew, harvest mouse and house mouse into ‘small mammals’; and the goldcrest, dunnock, magpie, great tit, jay, robin, skylark, chaffinch, wren, and thrush and sparrow species into ‘Passeriformes’. Some species were found in fewer than 5 % of scats even after grouping and were excluded from analysis - these were the large birds (chicken, curlew and Anatidae), fish (chub, grayling, and tilapia), and the common frog. A custom R script was used to perform all subsets variable selection based on the AICc, with gamebird release treatment and season included in every subset. Variable selection was run twice for each land use dataset. The interaction between season and gamebird release treatment was included as a potential covariate. Model fit was assessed via diagnostic plots (function plot.gllvm). The final model output was compared with a reduced model with only treatment and season as predictors to understand how the inclusion of land use measures affected the significance and estimates of the effect of treatment and season on affected taxa. Finally, the predict.gllvm function was used to predict the probability of species detection in each gamebird release treatment and season, if the significant land uses were the same across all sites (i.e. arable land cover and grass crop at their mean values across all sites).

## 3 Results

The libraries were sequenced on ∼1 % S4 flowcell which produces ∼10 billion reads, which resulted in a sequencing depth substantially higher than generally applied for studies of this nature. After all quality filtering, the mean number of reads per sample was 217,270 (SD = 155,479).

42 samples could not be confirmed to be from a fox. 12 samples had only fox reads, and a further 28 had only gamebird (once fox reads had been removed).

After quality filtering, 422 samples were deemed to have sufficient data for a robust analysis. More scats were found and therefore analysed from non-release sites (257 analysed) than from release sites (165 scats analysed). The number of scats found and analysed also differed substantially between seasons (Table 1).

**Table 1.** The number of scats from which suitable data was obtained and analysed, by release treatment and season. Number of sites from which the samples were taken is given in brackets.

|  | Post-shoot | Summer | Post-release | Winter | Total |
| --- | --- | --- | --- | --- | --- |
| **Release** | 59 (9) | 43 (10) | 50 (9) | 13 (4) | **165 (10)** |
| **Non-release** | 49 (8) | 113 (8) | 78 (7) | 17 (5) | **257 (8)** |
| **Total** | **108**<br>**(17)** | **156 (18)** | **128 (16)** | **30 (9)** |  |

### 3.1 Overall diet

The most frequently recorded vertebrate prey were mammals, which accounted for 74 % of observations. Rodents were the most highly predated mammals, with the field vole (*Microtus agrestis*) most frequently detected - accounting for 20 % of all observations, followed by the wood mouse or yellow-necked mouse (*Apodemus spp.*), the brown rat (*Rattus norvegicus*), the bank vole (*Myodes glareolus*), the Eastern grey squirrel (*Scuirus carolinensis*), and the Eurasian harvest mouse (*Micromys minutus*) (Figure 2). The common shrew (*Sorex araneus*) and pygmy shrew (*S. minutus*) together accounted for 0.5 % of observations only. The lagomorphs, including the European rabbit (*Oryctolagus cuniculus*) and brown hare (*Lepus europaeus*), accounted for ∼14 % of observations. The domestic dog (*Canis lupus*) accounted for ∼13 % of observations, making it the second most frequently detected species (likely resulting from coprophagy). A range of ungulates accounted for ∼ 6 % of observations, including roe deer (*Capreolus capreolus*), European fallow deer (*Dama dama*), cow (*Bos taurus*), sheep (*Ovis aries*), wild boar/domestic pig (*Sus scrofa*), and the Chinese muntjac (*Muntiacus reevesi*). Birds accounted for ∼25 % of observations, most of which were gamebirds (these data are presented in Sage et al., in prep.), followed by pigeons/doves (Columbidae), and broad diversity of passerine birds which accounted for ∼ 5 % of observations (Figure 2, Supplementary Table 1). Ground nesting bird species were rarely recorded with one scat containing curlew DNA and 5 scats (0.5 % of observations) contained skylark. Occasional detections of fish and amphibians accounted for < 1 % of observations (Figure 2, Supplementary Table 1).

**Figure 2.**
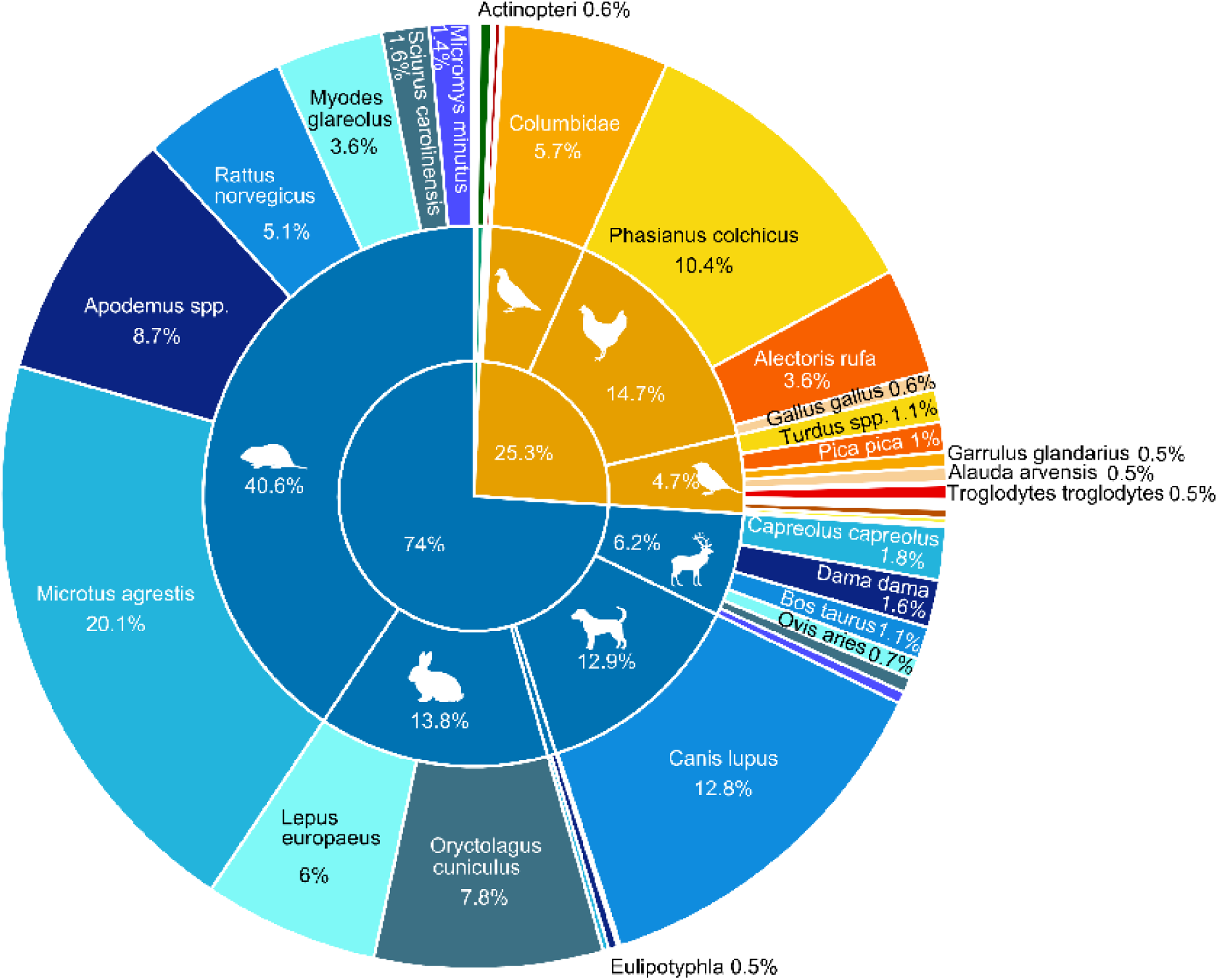
Percentage of observations of all taxa detected from all fox scat samples analysed. Values were calculated as the number of detections of a taxa / total detections across all taxa × 100. For visual clarity taxa < 0.5 % of observations have not been labelled. The numbers used to create the figure can be found in Supplementary Table 1.

### 3.2 Dietary diversity across gamebird release versus non-release sites and seasons

Observed richness, which counts all species regardless of abundance, and Hill-Shannon, which puts more weight on the common species, both varied significantly with gamebird release while neither were affected by crop types or land cover (Table 2). Both of these diversity metrics were significantly reduced on sites with gamebird release. For the Hill-Simpson metric, which lends the most weight to the dominant species, the crop type winter wheat was found to have a positive impact on diversity. A significant negative effect of gamebird release on Hill-Simpson diversity was only found when winter wheat was included in the model. The strength of effect of gamebird release was similar for all three metrics, but slightly lower for observed richness compared to Hill-Shannon and Hill-Simpson (Table 2).

**Table 2.**
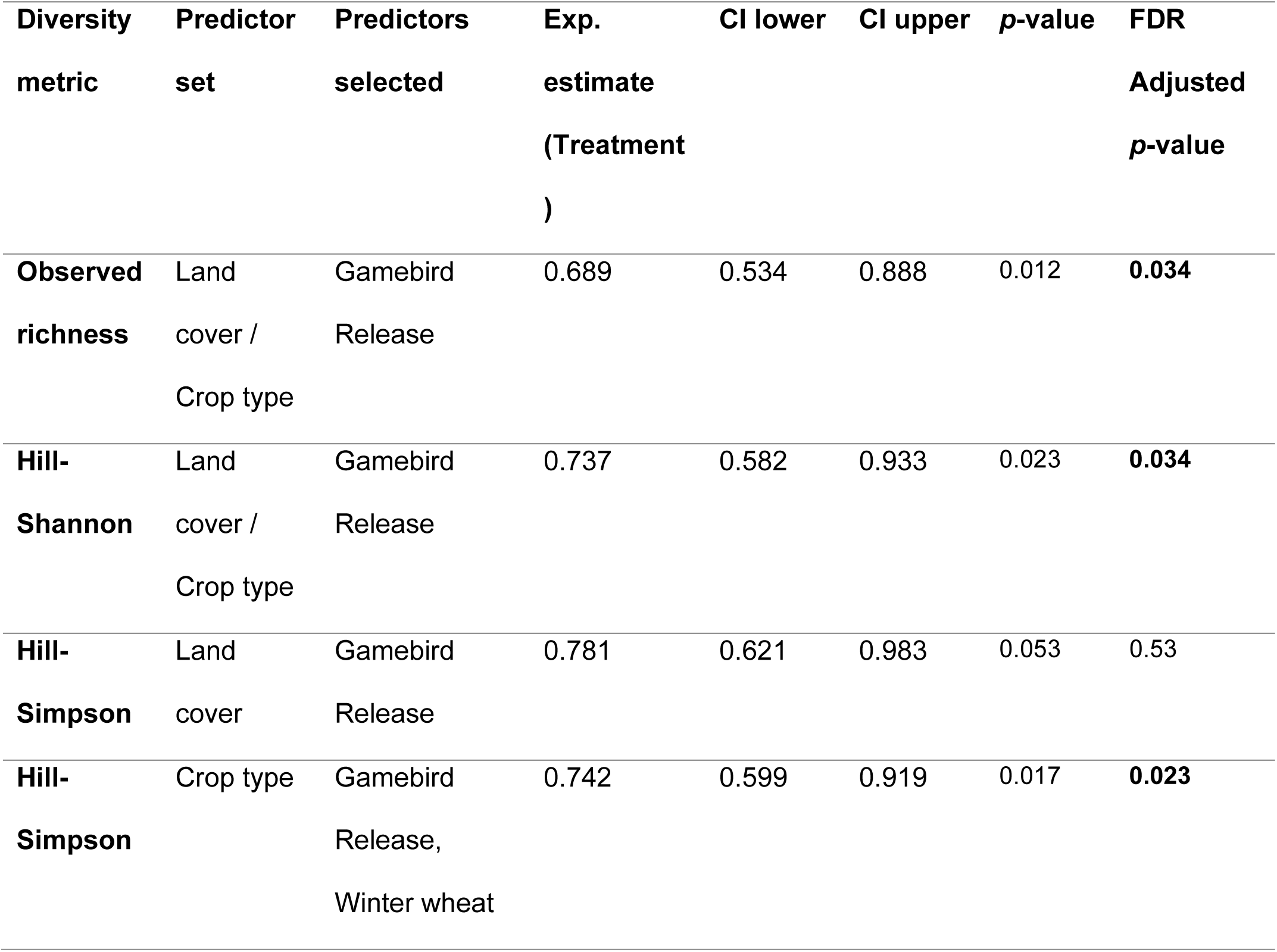
Results of model selection for the alpha diversity metrics. Estimates have been exponentiated to aid interpretation. Both raw p-values and p-values adjusted for multiple testing using the Benjamini–Hochberg false discovery rate (FDR) procedure are reported. One release site with only 3 scat samples was removed from the analysis, so the model included were 257 scat samples from 8 non-release sites and 162 scat samples from 9 release sites.

The lower dietary diversity on gamebird release compared to non-release sites was consistent throughout the year, but it was clearest in the post-shoot season (February to April) (Figure 3), when the release sites showed low variability in diversity across samples. In the other three seasonal periods, between sample variability was higher. On both site types, seasonal variation was observed, with a peak in diversity occurring in summer followed by a gradual reduction (Figure 3).

**Figure 3.**
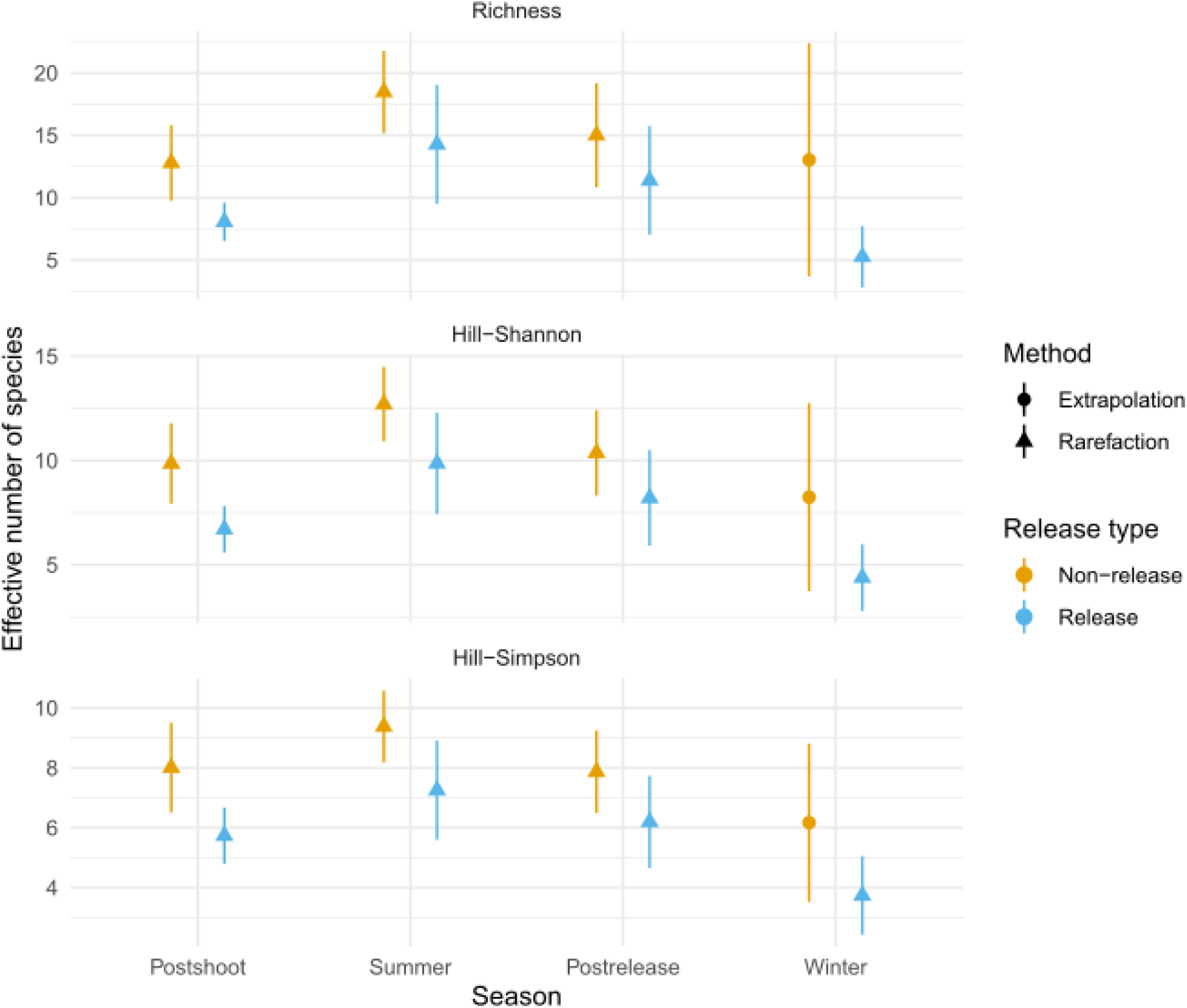
Comparison of species diversity metrics between gamebird release treatments and seasons. All samples from each treatment-season category were grouped together, thus does not account for individual sites. Sample coverage was standardised to 95 %. Seasons are defined as in Sage et al. (in prep.) – February to April are ‘post-shoot’, May to July are ‘summer’, August to October are ‘post-release’, and November to January are ‘winter’.

Figure 3 Comparison of species diversity metrics between gamebird release treatments and seasons. All samples from each treatment-season category were grouped together, thus does not account for individual sites. Sample coverage was standardised to 95 %. Seasons are defined as in Sage et al. (in prep.) – February to April are ‘post-shoot’, May to July are ‘summer’, August to October are ‘post-release’, and November to January are ‘winter’.

### 3.3 Dietary composition across gamebird release versus non-release sites and seasons

All subsets variable selection found the inclusion of percent cover of arable land and grass crop within 1 km to improve the model fit over the basic GLLMM with only gamebird release treatment and season as covariates. The crop type grass was not nested within arable land cover, so the two variables were considered independent and were included together in the final model.

Gamebird release was found to negatively affect the occurrence of domestic dog and bank vole, and positively affect the occurrence of the field vole, European rabbit and brown hare in the diet (Figure 4, Supplementary Table 2). While the grass crop and arable land cover estimates had significant effects on the occurrences of multiple taxa in the diet, an inspection of models with and without these covariates showed that the effects of gamebird release and season were consistent whether or not they were included. The exception to this was for the effect of gamebird release on the occurrence of the European rabbit. The cover of grass crop within 1 km was found to be associated with a significantly higher occurrence of the rabbit and grass crop occurred more on non-release sites (Figure 1). A positive effect of gamebird release on rabbit occurrence in the diet was only found when this effect of grass crop cover was accounted for in the model (Figure 4, Supplementary Table 2).

**Figure 4.**
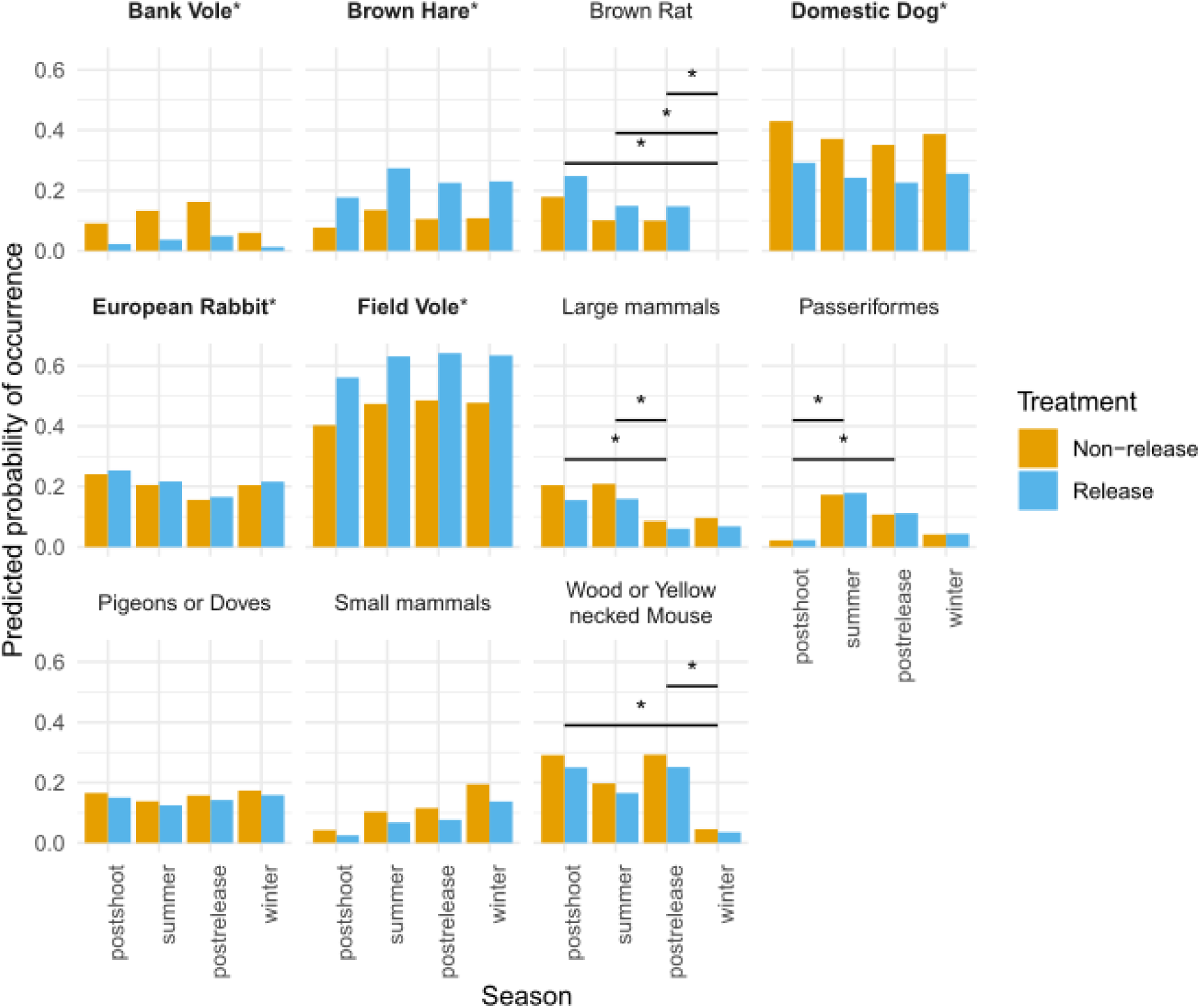
Variations in species probability of occurrence between gamebird release and non-release sites and across seasons, if crop types were equal between sites, as predicted by the GLMM. Species that differ significantly (p < 0.05) between gamebird release treatments are marked with an asterisk and bold name, significant differences between seasons are also marked with an asterisk.

Seasonal variations in the occurrence of taxa in the diet were also observed, which tended to follow the same pattern regardless of gamebird release treatment. Large mammals occurred more in the post-shoot and summer seasons, with occurrence declining in the post-release season. Passeriformes occurred more in the diet during the summer and post-release seasons than the post-shoot season. The wood / yellow-necked mouse occurred more in the post-shoot and post-release season than the winter. Finally, the brown rat was never detected in the winter and so occurred significantly more in all other seasons (Figure 4, Supplementary Table 2).

## Discussion

Applying DNA metabarcoding on fox scats collected across estates in Southern England, we identified a wide range of prey taxa to high taxonomic resolution that aligns generally well with previous studies for red fox diets more widely in the UK and across Europe. We also identified some key differences between diets in southern England and the rest of the UK and Europe, that likely reflect differences in prey availability and the methods of analysis used. We measured dietary diversity using three commonly used diversity metrics (observed richness, Hill-Shannon, and Hill-Simpson) that differ in the weight they give to rare and common species. All three metrics were significantly lower on gamebird release estates compared to non-release estates, which likely reflects changes in fox predatory behaviour caused by the abundance of gamebirds and/or variations in the availability of prey between the sites.

### Overall diet

Comparable studies from arable landscapes in Southern England have usually found rabbit to be the most common prey species (Baker et al., 2006; Reynolds et al., 1995; Reynolds & Tapper, 1995; Southern & Watson, 1941; Webbon et al., 2006), and small mammals such as field voles to be secondary prey. Here, we found the field vole to be the most frequently predated species. The only other study published reporting the field vole to be the primary prey species in England was conducted immediately following an outbreak of myxomatosis that substantially reduced the availability of rabbits (Lever, 1959). The published studies finding rabbit to be the most common were all conducted from 1980 to 2000 (Baker et al., 2006; Doncaster et al., 1990; Reynolds et al., 1995; Webbon et al., 2006) or prior to the arrival of myxomatosis in the UK (Southern & Watson, 1941). Between 1996 and 2021, there has been a 58 % decline in the rabbit population in England (Heywood et al., 2023), potentially due to the spread of rabbit haemorrhagic disease (Apha, 2020), and this may be at least part of the reason for the recent lower fox predation rates on rabbits. However, as the sites in our study were mostly arable, the lower predation rates on rabbits may also have resulted from rabbit control practiced by the estate management.

We also found a high frequency of domestic dog DNA in the fox scat, that was second only to the field vole, and which has rarely been previously reported. A likely reason for this difference is in the analytical methods used; most studies have used morphological analyses of undigested remains in the stomach and faeces, while we used a molecular approach to detect DNA. As there are very rarely dog carcasses available to scavenge in England, and we sampled the interiors of the scats to reduce environmental contamination, the source of this DNA is likely dog faeces which would not be detectable by morphology. This is supported by Waggershauser et al. (2022) who also found high occurrence of coprophagia of dog faeces from fox scats in Scotland using molecular methods. Furthermore, we consistently detected more dog DNA from non-release sites, suggesting foxes could be exploiting dog faeces, which are highly calorific, when prey is more scarce (Waggershauser et al., 2022). Alternatively, accessibility to the public and therefore the number of dogs passing through may have differed between the release and non-release site types, but we did not account for this.

We found little evidence of predation on ground nesting birds or other species of conservation concern, with 5 scats containing skylark and 1 containing curlew. Lapwing is the most common ground nesting farmland wader in the region (O’Brien & Smith, 1992) and was never detected in the scats, even before filtering low abundance OTUs. Preliminary work found that the primers used here amplify a wide range of birds with very little bias, so it is expected that DNA from ground nesting birds would be detected following predation on a chick or adult bird. However, it is possible that the metabarcoding assay used here is not sensitive enough to detect DNA from predation on bird eggs. During the earlier stages of development eggs contain very few cells, and therefore little DNA, which may not be detectable after having passed through the digestive system (Deagle et al., 2005; Thuo et al., 2019). It is therefore possible that the foxes did consume early-stage eggs.

### Variation in non-game diet with gamebird release

We found significantly lower dietary prey species diversity on release sites compared to non-release sites, particularly in the spring months following the shooting season. This difference was similar when measuring diversity with a range of metrics that weight rare and common species differently, indicating that the reduction in diversity was due to a reduction in predation frequency of both the common prey and the rare prey species.

It should be noted that the methods we employed excluded the possibility of detecting invertebrates in the diet, as the primers used were specific to vertebrates. We therefore cannot rule out the possibility that the variation in diversity may relate also to differences in the abundance and/or diversity of invertebrates on the sites. Some studies have found invertebrates to be an important component of the foxes’ diet; however, these were conducted in different environments than we have studied - pastureland, heathland, and forest (Williams et al., 2024), and urban (Doncaster et al., 1990). Studies from similar habitats to our sites (arable farmland in Southern England) have found that invertebrates tend to make up only a relatively small portion of the fox diet by weight/mass (Baker et al., 2006; Reynolds et al., 1995; Southern & Watson, 1941; Webbon et al., 2006) and the variation in invertebrate abundance would therefore have to be extreme to have an effect on the predation of vertebrates by foxes. It is therefore likely that the diversity difference observed is due to gamebird release and / or game management.

One particular concern regarding gamebird release is that the provision of easy prey or carrion could increase fox population density, and that once gamebird numbers have declined after the shooting season those foxes would switch to other prey species, causing decline in their populations (Madden & Sage, 2020; Mason et al., 2020). We found partial evidence to support this. On both release and non-release sites, the dietary diversity metrics followed a seasonal pattern – lower in the winter and post-shoot (spring) seasons, and highest in the summer, with diversity on the release sites always lower. This is in contrast with the pattern reported by Sage et al. (in prep.) on the seasonal frequency of gamebird in the foxes’ diet, where gamebird predation was highest on release sites and in the winter and post-shoot seasons. This contrasting pattern would be expected if the foxes switched to a more diverse prey spectrum following a decline in gamebird numbers. This is, however, correlational and does not confirm a cause and effect. This pattern could also be caused by the phenology of other prey species, such as the breeding season of passerine birds and large mammals which may increase their vulnerability in summer, or torpor in wood or yellow-necked mice which may decrease their availability to foxes.

Furthermore, the variation in dietary diversity was not solely due to gamebird predation, as other key prey species occurred more in the foxes’ diet on release sites, including during seasons when gamebird predation was high (Sage et al., in prep.). The lower overall dietary diversity on release sites is despite findings that gamebird release estates support high biodiversity, attributed to a mix of habitat improvements, supplementary feeding, and predator control (Madden et al., 2023; Madden & Sage, 2020; Mason et al., 2020; Sage et al., 2020). Gamebird release sites often support a high diversity of bird species (Draycott et al., 2008; Hoodless et al., 2005; Sage, 2018) as well as high abundances of multiple of the foxes’ preferred prey species. Brown hares and rodents in particular have been found to be positively associated with gamebird release (Coomes et al., 2023; Davey, 2008; Madden et al., 2023), which may explain the higher predation rates we observed on the field vole and the brown hare on release sites. The high availability of both gamebirds and other preferred prey species on release sites may allow the foxes to narrow their diet, thus reducing species diversity in their diet. This could protect other species from fox predation by diverting the predation pressure, and this may be part of the reason that gamebird release estates have been found to support high biodiversity. In addition, fewer fox scats were found on gamebird release estates indicative of lower fox activity (Sage et al., in prep.), likely due to effective fox control being carried out on release sites to protect the gamebirds. Reduced fox activity will mitigate the predation pressure on the key prey species and further protect the less frequently predated species.

Our finding of higher dietary diversity on non-release estates, combined with the finding that fox activity was higher on these sites (Sage et al., in prep), indicates higher predation pressure on a broad range of species. In particular, higher predation on bank voles was observed despite findings that they are positively associated with gamebird release (Davey, 2008). It could be that bank voles are less nutritious or more difficult to catch than the hare or field vole, but are preferred over passerine birds or shrews for example, and are therefore predated on only when more preferable species are less available. The bank vole and domestic dog (faeces, see above) were the only two species found significantly more on non-release sites and seem to be the primary food items the foxes’ switch to when more preferable prey is less available. The remaining dietary diversity found on non-release estates consisted of large mammals, passerine birds, pigeons and/or doves, wood/yellow-necked mice, and other small mammals. Although predation pressure by foxes appears to be lower for these species than the bank vole, the high fox activity could result in predation pressure that is limiting to some of these species’ abundances.

The effect of gamebird release on rabbit predation by foxes appears to be more complex and affected by gamebird release and management in more than one way. We found that there was a strong positive effect of grass crop on the occurrence of rabbit in the diet, with grass crop less common on release sites. A positive effect of gamebird release on rabbit predation by foxes was only found when grass crop cover was accounted for. The UKCEH ‘Land Cover plus: Crops 2022’ documentation does not provide clear definition of grass crop, but an almost perfect correlation (Pearson’s correlation coefficient = 0.96) between grass crop cover, and the sum of improved grassland and calcareous grassland on our sites suggests ‘grass crop’ refers to these two categories, with improved grassland being predominant on our sites. Improved grassland is characterised by the UKCEH as fast-growing grass species either used for pasture or regularly mown for silage or recreational purposes (*The UKCEH Land Cover Map for 2022*, 2024), resulting in short sward heights that are preferable to the European rabbit (Iason et al., 2002; Somers et al., 2012).

Gamebird estate managers often plant game cover crops to improve gamebird survival and to manage their movement (Henderson et al., 2004; Sage et al., 2005), which may be the reason for the reduced grassland on gamebird release sites. Thus, game management may be reducing the availability of rabbits to foxes via changes in crop plantations, and that another mechanism (e.g. supplementary feeding or predator control) is simultaneously increasing the availability of rabbits. This highlights the complex nature of the ecological effects of gamebird release and management.

The implications of these findings strongly depend on the fox population densities, and how this varies with gamebird release. Sage et al. (in prep.) found fewer fox scats on gamebird release estates compared to non-release estates, indicative of lower fox activity, most likely as a result of effective fox control by the estates’ management. This means lower predation pressure on all prey species, and more targeted predation on both gamebirds and particular species that are often more abundant on release sites will further protect the broader biodiversity on those sites. However, as suggested by Sage et al. (in prep.) and Sage et al. (2025), gamebird release may be supporting fox populations at a larger scale, as both gamebirds and foxes move in and out of gamebird release sites. The higher fox activity reported from the non-gamebird release estates, coupled with the high dietary diversity reported here, indicates strong and widespread predation pressure outside of gamebird release sites. Species such as the bank vole, which are predated on more where gamebirds are not released, may be particularly susceptible.

## Acknowledgements

The authors would like to express their gratitude to the British Association for Shooting and Conservation (BASC) and University of Exeter for co-funding this research and BASC for maintaining their strict separation from the research process. The project utilised sequencing equipment funded by the Wellcome Trust (Multi-User Equipment Grant award number 218247/Z/19/Z). For the purpose of open access, the author has applied a ‘Creative Commons Attribution (CC BY) licence to any Author Accepted Manuscript version arising from this submission.

## CRediT author statement

**Pedersen**: Conceptualization, Methodology, Formal analysis, Investigation, Data Curation, Visualization, Writing - Review & Editing, Writing - Original Draft. **Sage**: Conceptualization, Methodology, Investigation, Supervision, Resources, Writing - Review & Editing. **Woodburn**: Methodology, Investigation, Data Curation, Resources, Writing - Review & Editing. **Coomes**: Methodology, Investigation, Data Curation, Resources, Writing - Review & Editing. **Werling**: Investigation, Resources, Data Curation. **Tyler**: Conceptualization, Methodology, Resources, Supervision, Writing - Review & Editing

## Data availability

The raw sequencing data produced in this study can be found in the NCBI’s Sequence Read Archive under BioProject number PRJNA1379617. A matrix showing the presence/absence of each taxon in each sample can be found in Supplementary Table 2.

## Notes

### Competing Interest Statement

The authors have declared no competing interest.

